# Repulsive expansion dynamics in colony growth and patterning

**DOI:** 10.1101/2020.07.21.213611

**Authors:** Yangxiaolu Cao, John Neu, David R. Meldgin, Ting Lu, Lingchong You

**Author notes:** Corresponding author. Department of Biomedical Engineering, Duke University, CIEMAS 2355, 101 Science Drive, Box 3382, Durham, NC 27708, USA.

## Abstract

Spatial expansion of a population of cells can arise from growth of microorganisms, plant cells, and mammalian cells. It underlies normal or dysfunctional tissue development and it can be exploited as the foundation for programming spatial patterns. This expansion is often driven by continuous growth and division of cells within a colony, which in turn pushes the peripheral cells outward. This process generates a repulsion velocity field at each location within the colony. Here we show that this process can be approximated as coarse-grained repulsive-expansion kinetics. This framework enables accurate and efficient simulation of growth and gene expression dynamics in radially symmetric colonies with homogenous z-directional distribution. It is robust even if cells are not spherical and vary in size. The simplicity of the resulting mathematical framework also greatly facilitates generation of mechanistic insights.

## Introduction

Spatial expansion of a population of cells is ubiquitous in biology. It can arise from growth of bacterial or yeast colonies (1, 2), development of plant tissues (3), animal tissues (4), or growth of tumors (5–7). Understanding the expansion dynamics of a cell population is important, since colony expansion is a reflection of many cellular properties, including the substrate nutrient uptake rate, the metabolism, the division and death rates of individual cells. In addition, since colony expansion is usually coupled with the gene expression activity and the metabolic activity, understanding the expansion dynamics is the foundation of understanding the spatial dynamics of gene expression in growing colonies(8,9).

To date, primarily two types of models have been used to simulate spatial dynamics in growing colonies (including gene expression): agent-based models (ABMs) and continuum models based on partial differential equations (PDEs). An ABM focuses on simulating a single cell or a cluster of cells as an agent; the behavior of each agent is constrained by a set of pre-defined rules. The collective behavior of the population emerges from the interactions between agents, through contact or diffusible chemicals. Using the ABM entails assigning pre-defined rules and parameters to each agent. It provides detailed information about individual agents, as well as the overall system. However, it has several potential limitations. First, certain ABMs require more extensive assumptions about the definition of each agent, how each behaves, and how different agents interact (10)^-^(11). Many of these assumptions are made based on self-consistency and require parameters that are difficult or impossible to determine independently. Second, by construction, the computational demand by an ABM scales with the number of agents, as well as the complexity of reaction kinetics in each agent. As a result, ABMs are typically computationally expensive, making it difficult or even impossible to simulate the spatiotemporal dynamics of a cell population containing tens of millions of cells (a visible bacterial colony contains about 10-100 million cells). Finally, because of the sheer complexity of inter-agent interactions and the computational cost, it is difficult to deduce intuitive understanding of the properties emerging from the overall system (12).

These limitations can be partially alleviated by using continuum models consisting of PDEs. However, numerically solving PDEs can still be computationally demanding (though typically less so than ABMs) (13). Even though these tools can generate simulation results that can recapture certain aspects of the experimental results, it is difficult to develop intuitive understandings from the simulations.

In general, the colony expansion can be driven by a combination of internal and external forces. For instance, cells can preferentially move toward certain environmental cues, for example, by chemotaxis (14–16). They can also modulate motility by secreting and responding to surfactants, resulting in swarming (17–24).

In many cases, however, the expansion is primarily driven by continuous growth and division of cells in the colony (25). That is, the cells in the interior of a colony collectively push the peripheral cells outward, leading to colony expansion. The driver is the mechanical force generated by growth. This notion has been well recognized in diverse organisms, including bacteria (26, 27), yeast (28), plant cells (29, 30), and tumor cells (31, 32). Recognizing this feature enables us to model the colony expansion and gene expression using a highly simplified framework. In contrast to typical dimension-reduction of a complex model, the repulsive-expansion framework represents a completely different view in examining the spatiotemporal dynamics of expanding cell populations and associated gene expression. In particular, our model directly uses an ODE framework to track the “flow” of individual cells as they experience the steric forces generated due to cell growth and division. By reducing the structural complexity of the model, this framework facilitates development of mechanistic insights into the underlying dynamics of interest.

## Results

### The repulsive expansion model captures the colony growth dynamics

To develop our framework, we first assume the steric force between cell-cell growth and division are the driving force of colony expansion. At a particular location, the local steric force will generate a steric repulsion velocity field. We then assume that cells are spheres with the same radius. Therefore, on average, the overall pushing force on each cell is perpendicular to the boundary of a radial symmetry colony. If the profile of this velocity field is known with given initial conditions, one can use sets of ODEs to simulate the moving trajectories of any objects within the colony (Figure 1A).

**Figure 1:**
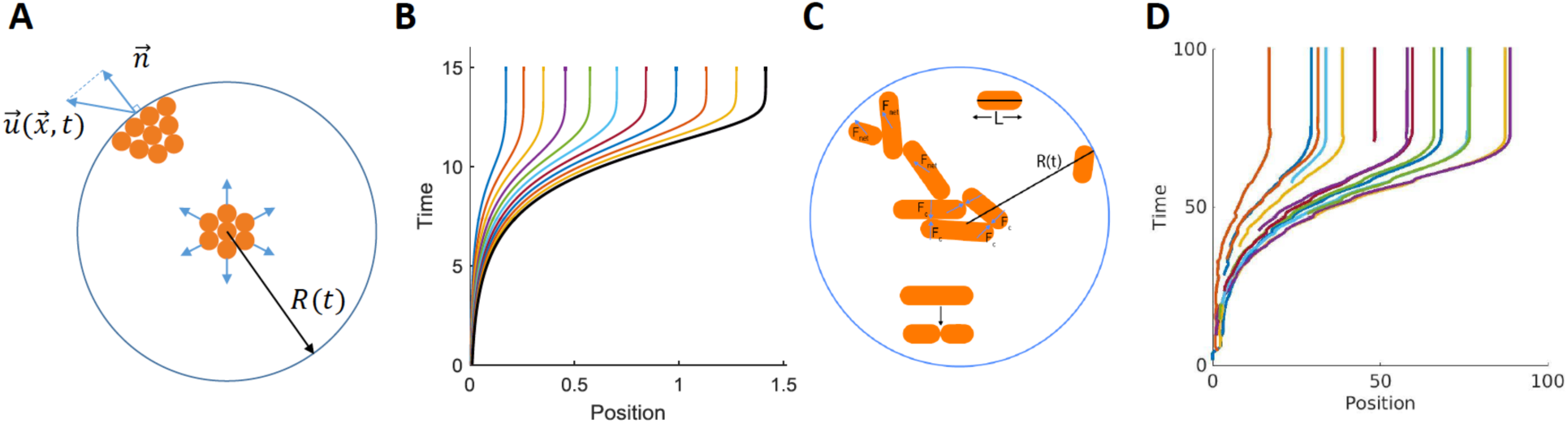
The repulsive expansion model captures colony growth dynamics. A. Illustration of the repulsive expansion dynamics. Each orange dot represents a cell at that given location. As cells in the colony grow and divide, cells in the interior will push the outer cells, generating a velocity field *u*(*x, t*). Given a location, 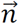 is the normal vector along radial direction. Therefore ***u*** *·* ***n*** represents the radial expansion velocity at the location. Since the model is assumed to be radial symmetric, the most outer layer of cells determines the size of the colony. At time *t*, the colony radius is expressed as *R*(*t*). B. Simulated colony growth over time using the repulsive expansion model. The black line indicates the radius of the colony, the colored lines represent the trajectories of pre-selected 50 locations within the colony. Each line corresponds to one particular initial position (*r*) within the colony. The outermost line correspond to the initial value of colony radius *R*. The initial r values can be set arbitrarily (*r* < *R*), without affecting the computational accuracy. C. Illustration of the ABM. Each orange rod represents a cell at a given location. Cells grow in length (*L*) by consuming nutrients. When a cell reaches a sufficient length, it divides into two shorter cells. Cell-cell contact interactions are modeled by elastic interactions. On average, contact forces and growth then produce a net force outward which drives colony expansion. The colony radius *R*(*t*) is determined by calculating the distance from the center to the furthest cell at time *t*. D. Simulated colony growth versus time by the ABM. Lines represent the radial distances of the 1^st^, 2^nd^, 4^th^, 48^th^, … 8192^th^ cells in time. Results qualitatively agree with those generated from the repulsive-expansion model.

In a closely packed colony, the cell division rate depends on the nutrient concentration and the location within the colony. That is, at the same nutrient concentration, if a cell is closer to the colony center, the slower the division rate is (33, 34). The rate of change in the colony volume can either be expressed as a function of the normal velocity of the colony boundary or the total number of the cells. In our framework, the velocity field of any given position (within a colony) is expressed as a function of cell division rate and the distance between the cellular location and the colony center (see Supplementary Material for detailed derivations). Knowing this velocity field, in turn, enables the computation of the movement of each cell.

Specifically, numerically solving the governing equations (Eq. 9, see Supplementary Table 1 for parameter values) can generate the dynamics of colony expansion as a function of time. Figure 1B shows a typical set of results on colony radius expansion (solid black line). If we choose a specific location within the colony and imagine there is a cell, we can plot how this cell moves based on the velocity field. Therefore, the moving trajectories of all cells in the colony can be tracked during the expansion (colored lines). This notion could be counter-intuitive if a cell does not exist initially and thus does not have a location in the initial colony. For example, consider a cell born at location *r*_1_ and at time *t*_1_. If a velocity field trajectory passes this position (*r*_1_, *t*_1_), the trajectory would predict how the cell would move at any time point later than *t*_1_. The section of the trajectory before *t*_1_ would be imaginary. However, to simplify presentation, all the trajectories are plotted from time zero in our simulation.

To evaluate the validity of the fundamental conceptual framework of the repulsive expansion model, we used an ABM (35) to simulate colony growth with similar physical constraints. Briefly, cells are modeled as elastic rods that increase in length as they consume nutrients. Cellular growth is modeled by a Monod equation (36) depending on nutrient availability. The total length of cell division is a truncated normal distribution, with values taken over a range of values (Supplementary Table 2). Repulsive growth is maintained by using a soft particle technique (37), which models cellular interactions as those between overlapping elastic spheres. ABMs were conducted to simulate the growth profiles of a single cell growing into 10,000 cells. The cell trajectories generated by an ABM simulation (Figure 1D) are qualitatively the same as those generated from the repulsive expansion model. In the ABM simulation, however, each trajectory only starts when a cell is born and does not have a hypothetic section leading to a position at time 0.

In the repulsive expansion model, we do not have cell shape or cell size as basic parameters. We used an ABM to validate that the colony expansion size will not be affected by cell shape or size (Figure 2A). While the repulsive expansion model is derived assuming spherical cells, it remains to be reliable even when this assumption is relaxed: the colony expansion dynamic from repulsive expansion model is comparable to the results solved from PDEs or ABM. For example, using repulsive expansion model (see Supplementary Table 3 for parameter values), regardless of the initial colony size, with the same environmental condition, the final colony sizes are the same (Figure 2B).

**Figure 2:**
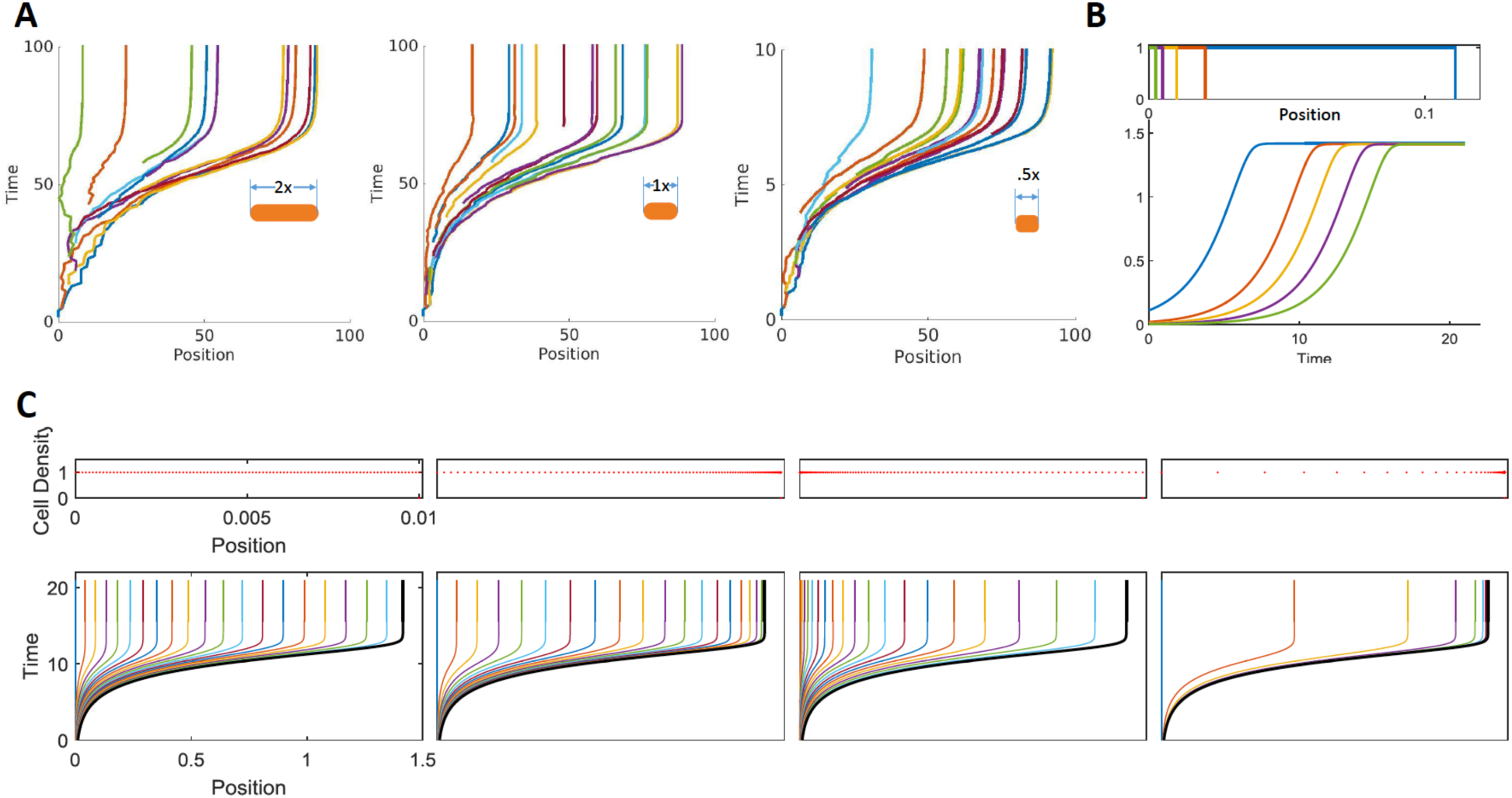
The repulsive expansion model under different initial model settings. A. Simulated colony growth using the ABM, assuming different cell aspect ratios. Colonies were grown with cells 2× (left), 1× (middle), and 0.5× (right) of the typically observed values for *E. coli.* The results demonstrate how changing cell size effects repulsive expansion. B. Simulated colony growth using the repulsive expansion model, assuming different initial colony sizes. The top panel shows four cell density distribution with the same shape, the same initial selected positions, but different initial colony sizes. The bottom panel shows the colony radius expansion over time with different initial colony sizes (same color code). C. Simulated colony growth using the repulsive expansion model, assuming different initial cell-density distributions. The top panel shows four cell density distributions with the same initial colony size and the same initial selected positions. The bottom panel shows the colony radius expansion over time with different cell density distributions (same color code). Thanks to the unique structure of the repulsive-expansion model, the accuracy of the computation is independent of how the initial positions are seeded for a given inoculum.

An important advantage of the repulsive expansion model is that the accuracy of computation does not depend on the resolution of the spatial discretization in initiating the simulation. This property drastically increases the computation efficiency without sacrificing computational accuracy. Despite differences in the segmentation of the spatial meshes, given the same initial colony size, the moving trajectories within the colony from different initial configurations are the same (Figure 2C). This property does not occur in the ABMs or PDE models, where a sufficiently high resolution in discretizing the space is critical to ensure computational accuracy or reliability. However, in the repulsive expansion model, a cell trajectory is directly determined by cell growth rate (Eq 9). Once the cell growth rate is given, the trajectory of each position could be expressed in an analytical solution, which does not require any spatial discretization. This feature will substantially simplify the simulation of cell movements.

### Modeling programmed pattern formation dynamics using the repulsive expansion model

The repulsive expansion model can be readily extended to describe gene expression dynamics that are coupled with colony expansion. To demonstrate this practice, we apply this framework to the analysis of a synthetic pattern-formation circuit that we recently engineered (38, 39). The circuit consists of a T7 RNA polymerase (T7) that activates its expression. Upon activation by T7, synthesis of AHL (A) will be mediated, which can diffuse across the cell membrane. When the global AHL concentration surpasses a threshold, intracellular AHL will trigger the activation of the synthesis of T7 lysozyme (L). Lysozyme then binds to the T7 and forms a T7-lysozyme complex (P), therefore inhibiting the T7 binding to the T7 promoter. This T7-lysozyme complex also inhibits T7 transcription. In this process, the AHL concentration is affected by its initial concentration and the domain size. The expression rates of T7, lysozyme, and AHL are all controlled by the spatially dependent gene expression capacity.

Depending on the experimental conditions, including cell strains and growth substrates, the circuit can generate different patterns. In particular, Payne et al. demonstrated the generation of one sharp ring or multiple rings when starting from single bacteria within a semi-solid agar droplet. Cao et al. demonstrated the generation of core-ring patterns that scale with the colony size when initiating the growth and patterning process from a few cells inkjet-printed to the top of agar surface. In both cases, the models are assumed to be radial symmetry. One study is in a one-dimensional ABM (39) and one used a PDE model (38).

Using comparable settings, the ABM was most time consuming – each single simulation took about 20hrs on our computer. The PDE model drastically improved the efficiency by reducing the time to ∼20min. With appropriate parameter choices, the PDE model can also reproduce the qualitative aspects of the outcomes from the ABM. The circuit dynamics can be readily implemented using the repulsive-expansion model (Eq10 in the Supplementary Material), which generates results comparable to the ABM (Figure 3B) or PDE (Figure 4B) models but with a speed that is 18,000-fold faster than the ABM model and 170-fold faster than the PDE model.

**Figure 3:**
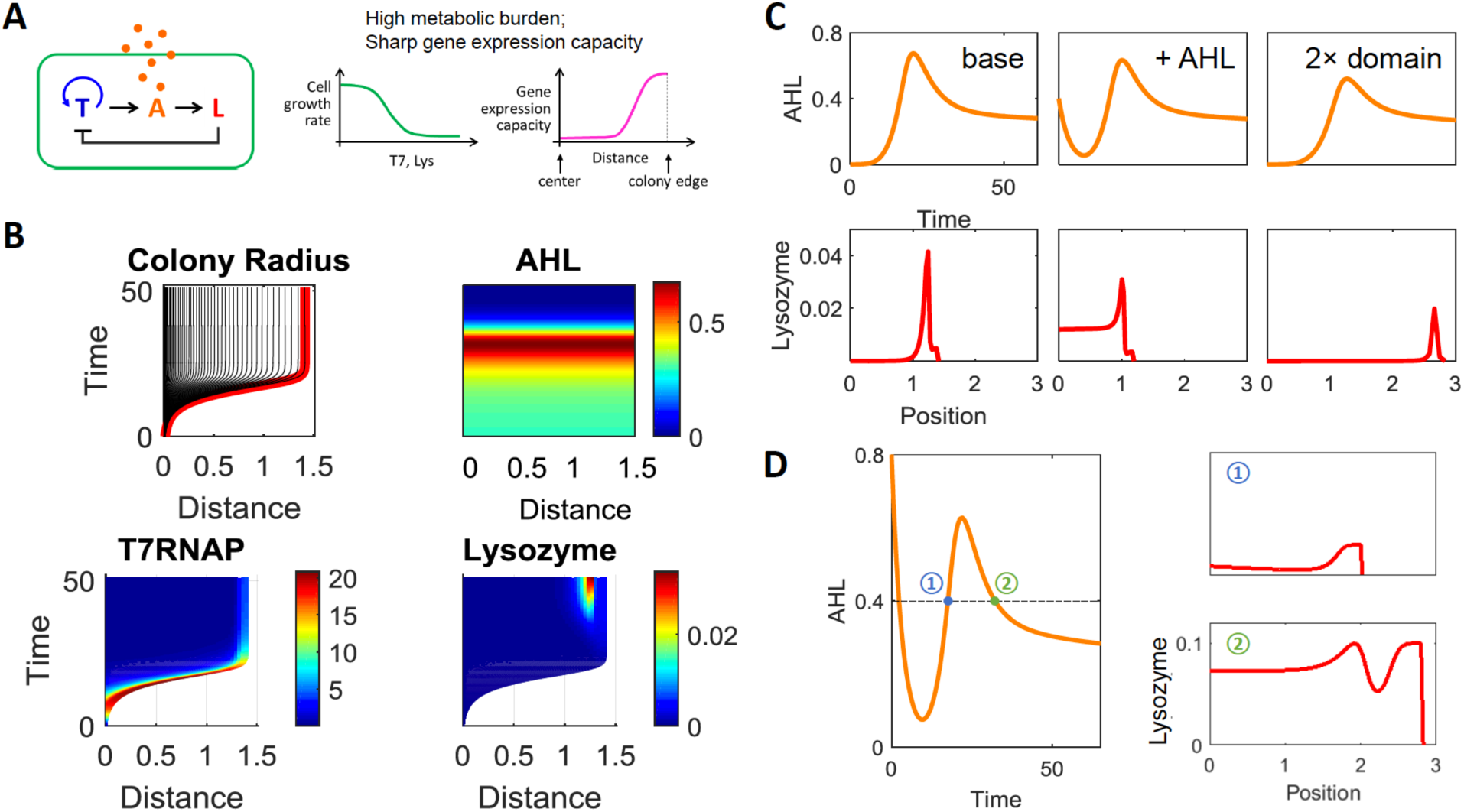
Simulated spatial-temporal dynamics of the pattern formation in engineered bacteria with a high metabolic burden and a sharp gene expression capacity profile. A. Left: circuit logic. Middle: the model was simulated under the condition of high metabolic burden. The cell growth rate is a decaying function of the production of T7 RNAP and T7 lysozyme. Right: Gene expression capacity. The x-axis represents the distance from the colony center. B. Top left to bottom right: Simulated spatial-temporal dynamics of colony radius, AHL, T7 RNAP and T7 lysozyme for varying distance (x-axis) over time (y-axis), respectively. The parameters used in the simulation are listed in Table 4. C. Simulated pattern formation with different environmental factors. The top panels from left to right shows the AHL dynamic under the base case (left, same with (B)), with initial AHL concentration is 0.3 (middle), and with domain size which is twice as large as the base case (right). The bottom panels are the T7 lysozyme distribution at the time when nutrient is exhausted under each condition, respectively. The x axis represents the distance from the colony edge. D. Simulated double rings. Left: the AHL dynamic with initial AHL concentration 0.8. Right top panel: the lysozyme distribution at time point 1, which is labeled in the left panel. Right bottom panel: the lysozyme distribution at time point 2.

**Figure 4:**
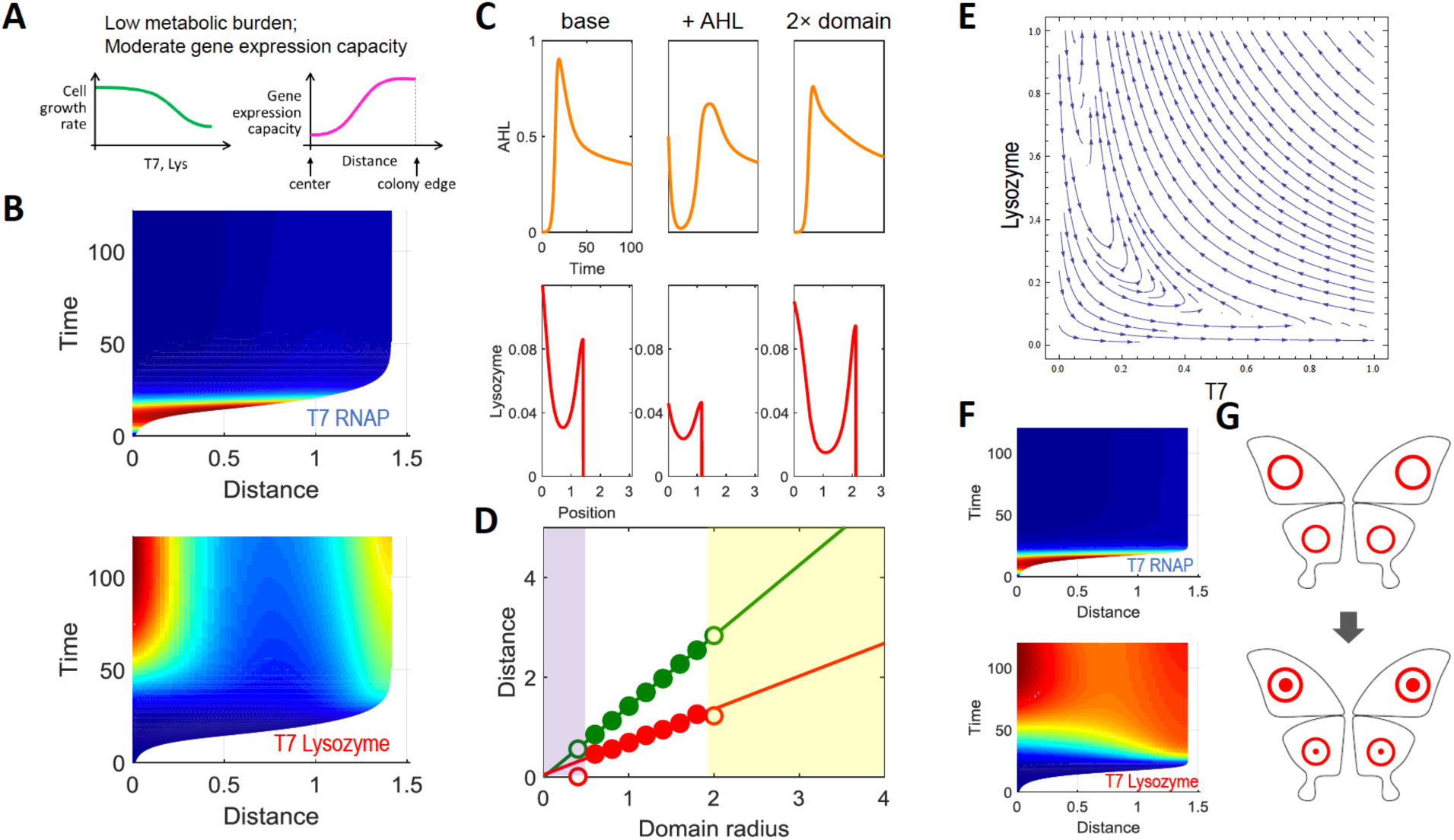
Simulated spatial-temporal dynamics of the pattern formation process with a high metabolic burden and a gene expression capacity profile with a moderate gradient. A. Simulation assumption with a low metabolic burden and a moderate gene expression capacity. The notations are the same as those in Figure 3. B. Simulated spatial-temporal dynamics of T7 RNAP and T7 lysozyme for varying distance (x-axis) over time (y-axis). The parameters used in the simulation are listed in Table 4, except *K*_*σ*_ = 0.1; *n*_*σ*_ = 2; *n* = 2; *K*_*φ*_ = 0.1. C. Simulated pattern formation with different environmental factors. The top panels from left to right shows the AHL dynamic under the base case (left), with initial AHL concentration 0.3 (middle), and with domain size which is twice as large as base case (right). The bottom panels are the T7 lysozyme distribution under each condition, respectively. The x-axis represents the distance from the colony edge. D. Simulated scale invariance in pattern formation. Dependence of the ring width (red circles) and the colony radius (green circles) on the domain radius from 1 to 4. The lines represent the linear regression of the colony radius and the ring width with respect to the domain radius in the white region. E. Phase diagram of T7 lysozyme (y axis) to T7 RNAP (x axis). The red line shows given an initial condition (T7RNAP=0.1, T7 lysozyme =0), T7 RNAP increases first then decreases, while T7 lysozyme keeps increasing. F. Simulated spatial-temporal dynamics of T7 RNAP and T7 lysozyme for varying distance (x-axis) over time (y-axis). The parameters used in the simulation are listed in the Table 4, except *K*_*σ*_ = 0.1; *n*_*σ*_ = 2. G. Ring pattern formation in the butterfly wings. Since eyespot signals to from inner and outer rings are released at different time-points. The outer rings form first, then the inner rings form within the outer rings at a later time point.

Similar to the analysis done in Cao et al, choosing appropriate parameters can allow us to generate the different patterns observed in both studies. When changing the environmental factors, the repulsive expansion model successfully recaptures the dynamics reported in Payne et al. Given an initial AHL concentration, a ring with a smaller radius forms around the colony edge compared to the one in the base case (without initial AHL). Given a larger initial domain size, a ring with a larger radius will form around the edge of a larger colony (Figure 3C). These results are consistent with the experimental and computational results presented in Payne et al. At the same domain size of the base case, two lysozyme rings can form given a high initial AHL concentration, which is consistent with the observations in Payne et al. (Figure 3D).

In the study by Cao et al., with an initial AHL concentration, a core and a ring with narrower width formed near the edge of the colony compared to the base case (without initial AHL). Given an initial larger domain, the repulsive expansion model indeed generates a wider core and ring will form in a larger colony (Figure 4C). Moreover, the simple model also successfully generates the scale invariance reported in Cao et al: the ring width and the colony radius are both proportional to the domain radius, when the latter is of moderate magnitude (Figure 4D).

In addition to the superior computational efficiency, the simplicity of the modeling framework provides more intuitive insights of circuit dynamics. For example, the dynamics of T7 is observed to be faster and more transient than lysozyme’s (Figure 4B). In Cao et al., the gene expression dynamics and the cellular movements are coupled in the PDEs. However, in the repulsive expansion model, particularly when the metabolic burden is small, the colony expansion could be modeled as independent of circuit dynamics. By separating the cellular movement from gene expression, we can simplify the model into two layers: the trajectories of all cells that reflect cellular movement, and the gene expression dynamic along each trajectory.

Since the repulsive expansion model is simpler and has fewer parameters, it provides a much clearer framework to reveal the essential properties of the system’s dynamics. By choosing an initial *r*, when *P* reaches equilibrium, we can draw the phase diagram of T7 and lysozyme. When T7 and lysozyme all start with small initial values, T7 will increase to a peak then decrease, while the significant accumulation of lysozyme happens after the peak of the T7 (Figure 4E). By combining these gene expression dynamics with the trajectories of all cells within the colony, we can see this correlation between the T7 patterns and lysozyme patterns. Compared with the ABM or the PDE model, the repulsive expansion model provides more intuition when it is used to analyze the dynamics of spatial patterns. In this case, the level of metabolic burden and gene expression capacity are two important factors to generate different types of patterns, which indicates the temporal cooperation between colony growth and gene expression are crucial for forming different types of patterns.

Since the repulsive expansion model describes cell moving trajectory and gene expression separately, this framework also provides direct insights into the relationship between cellular movement and circuit dynamics. In the previous PDE model, there are in total 22 parameters, which is very high dimensional. To reduce the complexity, the parameters that dictate to colony growth were fitted to experimental data. Cao et al. concluded the T7 lysozyme profile is mainly determined by circuit logic and growth dilution. During the colony expansion, near the colony edge, the lysozyme is insufficient to overcome the dilution of the cell growth. Therefore, a strong lysozyme core occurred before the ring formation. We agree the parameters being tested in the repulsive expansion model are a subset of the PDE and ABM’s parameters. Repulsive expansion model is not just a one-dimensional reduction of complicated models, but a model with core physic assumption so that redundant and trivial parameters can be eliminated for the researchers to quickly identify the key factors in affecting circuit dynamics.

The previous study has focused on the patterning process when the cell division and expansion rates are high. Given the simplicity of the repulsive expansion model, we can readily test the effects of the cell division and expansion rate when they are small. Interestingly, our new modeling result predicts the core-ring pattern formation with the ring forming before the core (Figure 4F). Lysozyme accumulates near the colony edge due to the high gene expression capacity at the location, which leads to ring formation. However, due to the circuit dynamic, the accumulating T7 and AHL lead to lysozyme increasing near the center of the colony. This patterning process of outer ring developing earlier than the inner core is analogous to the typical eyespot pattern-formation process in butterfly wings (40, 41). On the background of wings, there are a layer of parafocal elements (PFEs), which serve as the pattern units. The eyespot patterns are the PFEs with different color distributions. Many studies have shown the eyespot signals that form inner and outer rings are released at different time-points. The outer ring forms before the inner ring (Figure 4G) (42, 43). The network and the parameter combination identified by the repulsive expansion model could provide insights of how eyespot patterns are formed in butterfly wings.

## Discussion

In synthetic biology, tremendous progress has been made in programming single cells or cell populations by using gene circuits. The majority of these are done in a homogenous environment, e.g. in a liquid culture. Examples include generation of logic operations(44, 45), bistability (46, 47), and autonomous oscillations(48–50). Very few gene circuits have been successfully engineered to program self-organized spatial patterns(9, 38, 39, 51–53). The lack of progress in this direction is in major part due to the challenge associated with the design of gene circuits that could potentially generate desired function. This challenge, in turn, is due to the intrinsic difficulty in modeling spatial temporal dynamics of cell collectives involving cell growth and gene expression.

This difficulty is not unique to the design of synthetic gene circuits. Modeling analysis of colony expansion and patterning has a long history in endeavors to understand growth of bacterial physiology(1, 2), biofilm formation(54, 55), and tumor expansion(5). In general, spatiotemporal dynamics of colony expansion and gene expression (of signaling) are modeled by using ABMs or reaction-diffusion models consisting of PDEs. Both modeling frameworks suffer from two major limitations: high computational cost and difficulty in drawing intuition. Extensive ABMs have been developed to simulate tight-packed expanding colonies consisting of no motile bacteria (56–58). A one-dimensional molecular dynamic model was used to compare with the ABM model in one of the study (56), by directly accounting for the elastic and friction forces experienced by the cells. Our repulsive-expansion model represents another layer of simplification by focusing on the velocity field as a reflection of contact-based forces between cells.

Our work demonstrates a highly simplified yet accurate modeling framework for a particular class of dynamics involving colony growth and patterning. In essence, the repulsive expansion framework models the “flow” of cells in a growing colony, driving by the force generated by collective growth and division of cells in the colony. One can imagine a perfect radial symmetric colony, given the nutrient concentration and distribution, the colony growth rate of each cell at any position is known. Therefore, the trajectory of any cell with a given initial position can be calculated based on the force the cell experiences. This framework is structurally different from the corresponding PDE model and it does not arise from typical dimensional reduction. Due to its structural simplicity, this modeling framework offers superior computational efficiency as it does not require high resolution of space discretization to ensure computational accuracy. The framework is readily amendable to the modeling of spatial temporal dynamics coupled with or arising from colony growth and expansion. We have illustrated this point by applying the modeling framework to a synthetic gene circuit that has been previously analyzed.

In general, the framework could be applied to a class of spatial distribution problems: the system is radially symmetric, with homogenous z-directional distribution, and with gradient-free chemicals or chemicals that have gradient but exceeds the triggering threshold. These criteria are the cornerstone of the repulsive expansion dynamics. Many examples of biological processes indeed satisfy these conditions. Examples include colony growth and gene expression in a microfluidic chip with a confined chamber height (59–61), a single layer of cell divisions in a microscope slide (62–64), and cells in 3D symmetric spherical growth condition (65, 66). Even if these systems can secrete diffusive chemicals as global signaling molecules, our modeling framework is applicable if signaling molecules’ diffusion rates are sufficiently fast to form uniform spatial distributions (38, 39), or if the molecules accumulate fast enough to exceeds beyond the gene expression triggering threshold (67, 68).

## Author contributions

LY, JN, and YC conceived the project. JN proposed and derived the equations. YC developed numerical simulations in MATLAB. DRM and TL developed ABM simulations. YC, JN, and LY wrote the manuscript, with inputs from DRM and TL.

## Acknowledgements

This study was partially supported by the Office of Naval Research (LY), National Science Foundation (L.Y.), and a David and Lucile Packard Fellowship (L.Y.).

## References

1. Weijer CJ (2009) Collective cell migration in development. Journal of cell science 122(Pt 18):3215–23.

2. Zhang HP, Be’er A, Florin E-L, Swinney HL (2010) Collective motion and density fluctuations in bacterial colonies. Proceedings of the National Academy of Sciences of the United States of America 107(31):13626–30.

3. Milthorpe FL, Newton P (1963) Studies on the expansion of the leaf surface: III. The influence of radiation on cell division and leaf expansion. Journal of Experimental Botany 14(3):483–495.

4. O’Brien LE, Zegers MMP, Mostov KE (2002) Building epithelial architecture: Insights from three-dimensional culture models. Nature Reviews Molecular Cell Biology 3(7):531–537.

5. Friedl P, Hegerfeldt Y, Tusch M (2004) Collective cell migration in morphogenesis and cancer. International Journal of Developmental Biology 48(5–6):441–449.

6. Byrne HM (1999) A weakly nonlinear analysis of a model of avascular solid tumour growth. Journal of mathematical biology 39(1):59–89.

7. Greenspan HP (1976) On the growth and stability of cell cultures and solid tumors. Journal of Theoretical Biology 56(1):229–242.

8. Mendelson NH, Salhi B (1996) Patterns of reporter gene expression in the phase diagram of Bacillus subtilis colony forms. Journal of Bacteriology. doi: 10.1128/jb.178.7.1980-1989.1996.

9. Liu J, et al. (2015) Metabolic co-dependence gives rise to collective oscillations within biofilms. Nature 523(7562):550–554.

10. Railsback SF, Lytinen SL, Jackson SK (2006) Agent-based Simulation Platforms: Review and Development Recommendations. Simulation 82(9):609–623.

11. Morvan G (2012) Multi-level agent-based modeling - Bibliography. Foundations:1–13.

12. Gorochowski TE (2016) Agent-based modelling in synthetic biology. Essays In Biochemistry. doi: 10.1042/EBC20160037.

13. Haberman R (2015) Applied Partial Differential Equations doi: 10.1007/978-0-387-74372-1.

14. Henrichsen J (1972) Bacterial surface translocation: a survey and a classification. Bacteriological reviews 36(4):478–503.

15. Harshey RM (2003) Bacterial Motility on a Surface: Many Ways to a Common Goal. Annual Review of Microbiology 57(1):249–273.

16. Verstraeten N, et al. (2008) Living on a surface: swarming and biofilm formation. Trends in Microbiology 16(10):496–506.

17. Kearns DB (2010) A field guide to bacterial swarming motility. Nature Reviews Microbiology 8(9):634–644.

18. Mobley HLT, Belas R (1995) Swarming and pathogenicity of Proteus mirabilis in the urinary tract. Trends in Microbiology 3(7):280–284.

19. McCarter L, Silverman M (1990) Surface???induced swarmer cell differentiation of Vibrio parahaemoiyticus. Molecular Microbiology 4(7):1057–1062.

20. Kearns DB, Losick R (2003) Swarming motility in undomesticated Bacillus subtilis. Molecular Microbiology 49(3):581–590.

21. Harshey RM (1994) Bees aren’t the only ones: swarming in Gram-negative bacteria. Molecular Microbiology 13(3):389–394.

22. Eberl L, Molin S, Givskov M (1999) Surface motility of Serratia liquefaciens MG1. Journal of Bacteriology 181(6):1703–1712.

23. Hall PG, Krieg NR (1983) Swarming of *Azospirillum brasilense* on solid media. Canadian Journal of Microbiology 29(11):1592–1594.

24. Hernández F, Rodríguez E (1993) The swarming phenomenon of Clostridium tetani. Revista de biologia tropical 41(3B):857–9.

25. Farrell FDC, Hallatschek O, Marenduzzo D, Waclaw B (2013) Mechanically driven growth of quasi-two-dimensional microbial colonies. Physical Review Letters. doi: 10.1103/PhysRevLett.111.168101.

26. Farrell FD, Gralka M, Hallatschek O, Waclaw B (2017) Mechanical interactions in bacterial colonies and the surfing probability of beneficial mutations. Journal of the Royal Society, Interface 14(131):20170073.

27. Su PT, et al. (2012) Bacterial Colony from Two-Dimensional Division to Three-Dimensional Development. PLoS ONE 7(11). doi: 10.1371/journal.pone.0048098.

28. Kayser J, Schreck CF, Yu Q, Gralka M, Hallatschek O (2018) Emergence of evolutionary driving forces in pattern-forming microbial populations. Philosophical transactions of the Royal Society of London Series B, Biological sciences 373(1747):20170106.

29. Hamant O, Haswell ES (2017) Life behind the wall: sensing mechanical cues in plants. BMC Biology 15(1):59.

30. Dupuy L, Mackenzie J, Haseloff J (2010) Coordination of plant cell division and expansion in a simple morphogenetic system. Proceedings of the National Academy of Sciences 107(6):2711–2716.

31. Katira P, Bonnecaze RT, Zaman MH (2013) Modeling the Mechanics of Cancer: Effect of Changes in Cellular and Extra-Cellular Mechanical Properties. Frontiers in Oncology 3:145.

32. Mammoto T, Ingber DE (2010) Mechanical control of tissue and organ development. Development (Cambridge, England) 137(9):1407–20.

33. Varon M, Choder M (2000) Organization and cell-cell interaction in starved Saccharomyces cerevisiae colonies. Journal of Bacteriology. doi: 10.1128/JB.182.13.3877-3880.2000.

34. Shao X, et al. (2017) Growth of bacteria in 3-d colonies. PLoS Computational Biology. doi: 10.1371/journal.pcbi.1005679.

35. Blanchard AE, Lu T (2015) Bacterial social interactions drive the emergence of differential spatial colony structures. BMC Systems Biology. doi: 10.1186/s12918-015-0188-5.

36. Monod J (1949) The Growth of Bacterial Cultures. Annual Review of Microbiology. doi: 10.1146/annurev.mi.03.100149.002103.

37. Volfson D, Kudrolli A, Tsimring LS (2004) Anisotropy-driven dynamics in vibrated granular rods. Physical Review E - Statistical Physics, Plasmas, Fluids, and Related Interdisciplinary Topics. doi: 10.1103/PhysRevE.70.051312.

38. Cao Y, et al. (2016) Collective Space-Sensing Coordinates Pattern Scaling in Engineered Bacteria. Cell 165(3):620–630.

39. Payne S, et al. (2013) Temporal control of self-organized pattern formation without morphogen gradients in bacteria. Molecular Systems Biology 9(1). doi: 10.1038/msb.2013.55.

40. Otaki JM (2012) Structural analysis of eyespots: Dynamics of morphogenic signals that govern elemental positions in butterfly wings. BMC Systems Biology. doi: 10.1186/1752-0509-6-17.

41. Otaki JM (2017) Self-similarity, distortion waves, and the essence of morphogenesis: A generalized view of color pattern formation in butterfly wings. Diversity and Evolution of Butterfly Wing Patterns: An Integrative Approach doi: 10.1007/978-981-10-4956-9_7.

42. Iwata M, Ohno Y, Otaki JM (2014) Real-time in vivo imaging of butterfly wing development: Revealing the cellular dynamics of the pupal wing tissue. PLoS ONE. doi: 10.1371/journal.pone.0089500.

43. Otaki JM (2011) Generation of Butterfly Wing Eyespot Patterns: A Model for Morphological Determination of Eyespot and Parafocal Element. Zoological Science. doi: 10.2108/zsj.28.817.

44. Tamsir A, Tabor JJ, Voigt CA (2011) Robust multicellular computing using genetically encoded NOR gates and chemical “wiresg.” Nature. doi: 10.1038/nature09565.

45. Moon TS, Lou C, Tamsir A, Stanton BC, Voigt CA (2012) Genetic programs constructed from layered logic gates in single cells. Nature. doi: 10.1038/nature11516.

46. Gupta C, López JM, Ott W, Josić K, Bennett MR (2013) Transcriptional delay stabilizes bistable gene networks. Physical Review Letters. doi: 10.1103/PhysRevLett.111.058104.

47. Bennett MR, Hasty J (2007) A DNA methylation-based switch generates bistable gene expression. Nature Genetics. doi: 10.1038/ng0207-146.

48. Stricker J, et al. (2008) A fast, robust and tunable synthetic gene oscillator. Nature. doi: 10.1038/nature07389.

49. Hasty J, Dolnik M, Rottschäfer V, Collins JJ (2002) Synthetic gene network for entraining and amplifying cellular oscillations. Physical Review Letters. doi: 10.1103/PhysRevLett.88.148101.

50. Bratsun D, Volfson D, Tsimring LS, Hasty J (2005) Delay-induced stochastic oscillations in gene regulation. Proceedings of the National Academy of Sciences. doi: 10.1073/pnas.0503858102.

51. Karig D, et al. (2018) Stochastic Turing patterns in a synthetic bacterial population. Proceedings of the National Academy of Sciences. doi: 10.1073/pnas.1720770115.

52. Liu C, et al. (2011) Sequential establishment of stripe patterns in an expanding cell population. Science. doi: 10.1126/science.1209042.

53. Liu J, et al. (2017) Coupling between distant biofilms and emergence of nutrient time-sharing. Science. doi: 10.1126/science.aah4204.

54. Battin TJ, et al. (2007) Microbial landscapes: New paths to biofilm research. Nature Reviews Microbiology. doi: 10.1038/nrmicro1556.

55. Noguera DR, Okabe S, Picioreanu C (1999) Biofilm modeling: Present status and future directions. Water Science and Technology. doi: 10.1016/S0273-1223(99)00178-X.

56. Volfson D, Cookson S, Hasty J, Tsimring LS (2008) Biomechanical ordering of dense cell populations. Proceedings of the National Academy of Sciences. doi: 10.1073/pnas.0706805105.

57. Boyer D, et al. (2011) Buckling instability in ordered bacterial colonies. Physical Biology. doi: 10.1088/1478-3975/8/2/026008.

58. Mather W, Mondragón-Palomino O, Danino T, Hasty J, Tsimring LS (2010) Streaming instability in growing cell populations. Physical Review Letters. doi: 10.1103/PhysRevLett.104.208101.

59. Groisman A, et al. (2005) A microfluidic chemostat for experiments with bacterial and yeast cells. Nature Methods 2(9):685–689.

60. Jang K-J, Suh K-Y (2010) A multi-layer microfluidic device for efficient culture and analysis of renal tubular cells. Lab on a Chip. doi: 10.1039/B907515A.

61. Lopatkin AJ, et al. (2016) Antibiotics as a selective driver for conjugation dynamics. Nature Microbiology. doi: 10.1038/nmicrobiol.2016.44.

62. Elowitz MB, Leibier S (2000) A synthetic oscillatory network of transcriptional regulators. Nature 403(6767):335–338.

63. Mihalcescu I, Hsing W, Leibler S (2004) Resilient circadian oscillator revealed in individual cyanobacteria. Nature. doi: 10.1038/nature02533.

64. Tan C, Marguet P, You L (2009) Emergent bistability by a growth-modulating positive feedback circuit. Nature Chemical Biology. doi: 10.1038/nchembio.218.

65. Discher DE, Mooney DJ, Zandstra PW (2009) Growth factors, matrices, and forces combine and control stem cells. Science 324(5935):1673–1677.

66. Shirinifard A, et al. (2009) 3D multi-cell simulation of tumor growth and angiogenesis. PLoS ONE. doi: 10.1371/journal.pone.0007190.

67. Chen H, Xu Z, Mei C, Yu D, Small S (2012) A system of repressor gradients spatially organizes the boundaries of bicoid-dependent target genes. Cell. doi: 10.1016/j.cell.2012.03.018.

68. Roth S, Lynch J (2012) Does the bicoid gradient matter? Cell. doi: 10.1016/j.cell.2012.04.007.

